# Inferring parameters of cancer evolution from sequencing and clinical data

**DOI:** 10.1101/2020.11.18.387837

**Authors:** Nathan Lee, Ivana Bozic

## Abstract

As a cancer develops, its cells accrue new mutations, resulting in a heterogeneous, complex genomic profile. We make use of this heterogeneity to derive simple, analytic estimates of parameters driving carcinogenesis and reconstruct the timeline of selective events following initiation of an individual cancer. Using stochastic computer simulations of cancer growth, we show that we can accurately estimate mutation rate, time before and after a driver event occurred, and growth rates of both initiated cancer cells and subsequently appearing subclones. We demonstrate that in order to obtain accurate estimates of mutation rate and timing of events, observed mutation counts should be corrected to account for clonal mutations that occurred after the founding of the tumor, as well as sequencing coverage. We apply our methodology to reconstruct the individual evolutionary histories of chronic lymphocytic leukemia patients, finding that the parental leukemic clone typically appears within the first fifteen years of life.

## Introduction

When a cell accrues a sequence of driver mutations – genetic alterations that provide a proliferative advantage relative to surrounding cells – it can begin to divide uncontrollably and eventually develop the complex features of a cancer [1–3]. Thousands of specific driver mutations have been implicated in carcinogenesis, with individual tumors harboring from few to dozens of drivers, depending on the cancer type [4]. Mutations that don’t have a significant effect on cellular fitness also arise, both before and after tumor initiation [5]. These neutral mutations, or “passengers”, can reach detectable frequencies by random genetic drift or the positive selection of a driver mutation in the same cell [6–9]. Mutational burden detectable by bulk sequencing reveals tens to thousands of passengers per tumor [10, 11].

Genome sequencing technologies have revealed the heterogeneous, informative genetic profiles produced by the evolutionary process driving carcinogenesis [12, 13]. These genetic profiles have been used to obtain insight into specific features of the carcinogenic process operating in individual patients. For example, the molecular clock feature of passenger mutations has been employed to measure timing of early events in tumor formation, as well as identify stages of tumorigenesis and metastasis [14–22]. Other studies have estimated mutation rates [5, 23, 24], selective growth advantages of cancer subclones [25–28], and the effect of spatial structure on cancer evolution [29–31]. We note that previous approaches typically only estimate one or a few parameters of cancer evolution. In addition, many state of the art methods make use of computationally expensive approaches [24, 30, 32] or simplifying assumptions, such as approximating tumor expansion as deterministic or ignoring cell death [27, 32].

Mathematical models of cancer progression, especially when used in conjunction with experimental and clinical data, can provide important insights into the evolutionary history of cancer [9, 19, 33–37]. Branching processes – a type of a stochastic process – can be used to model how different populations of dividing, dying, and mutating cells in a tumor evolve over time [38]. Their theory and applications have been well developed to model the multistage nature of cancer development [25, 29, 35, 38–40]. Here we use a branching process model of carcinogenesis to derive a comprehensive reconstruction of an individual tumor’s evolution.

Tumors can grow for many years, even decades, before they reach detectable size [16]. Typically, tumor samples used for sequencing would be obtained at the end of the tumor’s natural, untreated progression. More recently, longitudinal sequencing, where a tumor is sequenced at multiple times during its development, has provided better resolution of tumor growth dynamics and evolution in various cancer types [27, 41– 44]. We establish that two longitudinal bulk sequencing and tumor size measurements are sufficient to reconstruct virtually all parameters (mutation rate, growth rates, times of appearance of driver mutations, and time since the driver mutation) of cancer evolution in individual patients. Our analytic approach yields simple formulas for the parameters; thus estimation of the parameters governing cancer growth is not computationally intensive, regardless of tumor size. Our framework makes possible a personalized, high-resolution reconstruction of a tumor’s timeline of selective events and quantitative characterization of the evolutionary dynamics of the subclones making up the tumor.

## Results

### Model

We consider a multi-type branching process of tumor expansion (Fig. 1a). Tumor growth is started with a single initiated cell at time 0. Initiated tumor cells divide with rate *b* and die with rate *d*. These cells already have the driver mutations necessary for expansion, so we assume *b > d*. The population of initiated cells can go extinct due to stochastic fluctuations, or survive stochastic drift and start growing (on average) exponentially with net growth rate *r* = *b*−*d*. We will focus only on those populations that survived stochastic drift.

**Figure 1:**
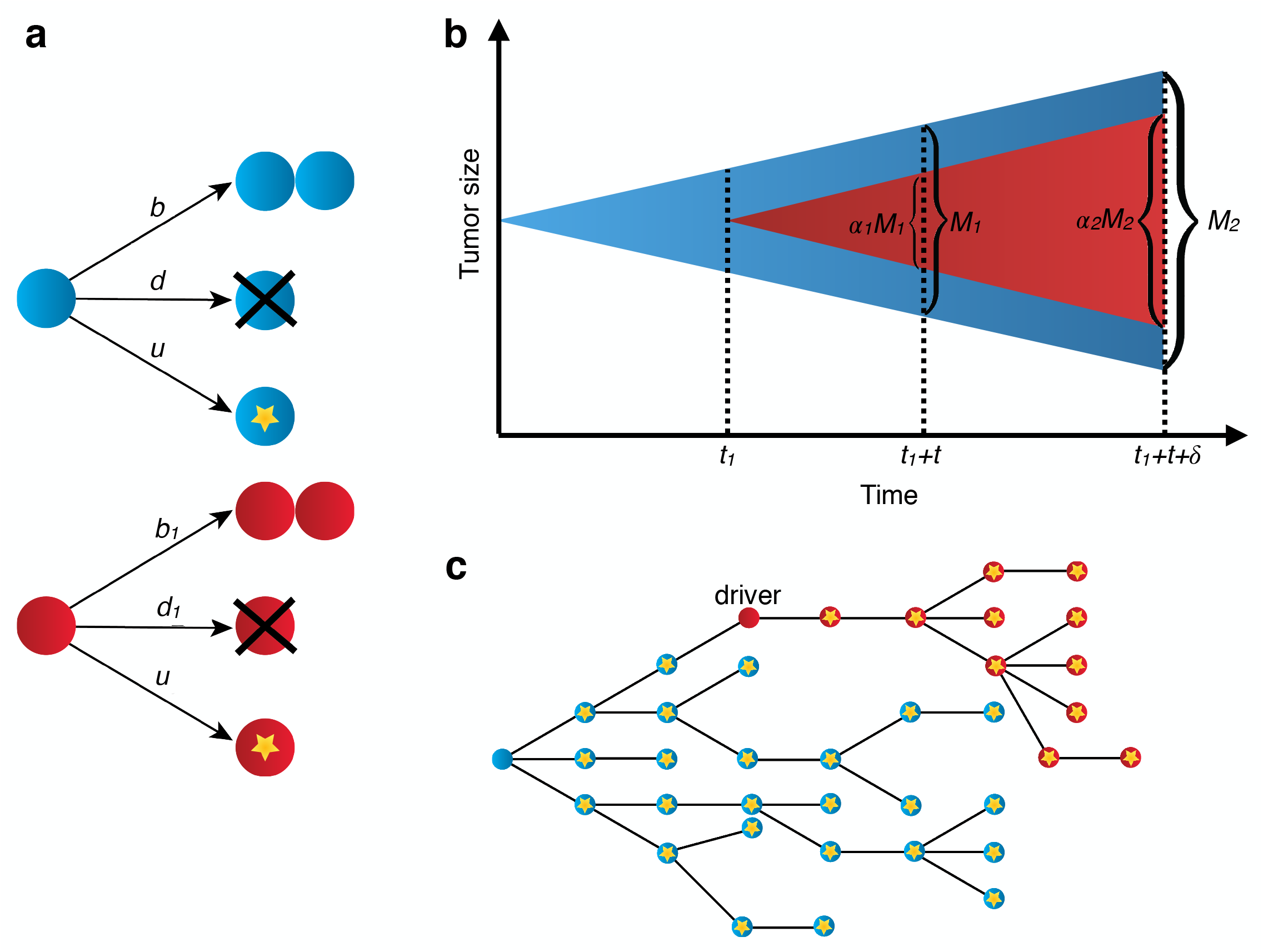
Stochastic branching process model of tumor evolution. (a) Stochastic branching process model for tumor expansion. Initiated tumor cells (blue) divide with birth rate *b*, die with death rate *d*, and accrue passenger mutations with mutation rate *u*. Type-1 cells, which carry the driver mutation, divide with birth rate *b*_1_, die with death rate *d*_1_, and accrue passenger mutations with mutation rate *u*. (b) The initiated tumor, or type-0, (blue) population growth is initiated from a single cell. A driver mutation occurs in a single type-0 cell at time *t*_1_, starting the type-1 population (red). The tumor is bulk sequenced at times *t*_1_ + *t* and *t*_1_ + *t* + *δ*. (c) By the time the tumor is observed, it has a high level of genetic heterogeneity due to the mutations that have accrued in both type-0 (blue) and type-1 populations (red). Each yellow star represents a different passenger mutation.

At some time *t*_1_ *>* 0 a new driver mutation occurs in a single initiated tumor cell, starting a new independent birth-death process, with birth rate *b*_1_ and death rate *d*_1_ (Fig. 1b). Net growth rate of cells with the new driver is *r*_1_ = *b*_1_ − *d*_1_. The new driver increases the rate of growth, i.e., *r*_1_ *> r*. We define the driver’s selective growth advantage by *g* = (*r*_1_*/r* − 1). In addition, both populations of cells (with and without the driver) accrue passenger mutations with rate *u* (Fig. 1c).

After the driver mutation occurs, an additional time *t* passes before the tumor is observed. Cells containing *i* new driver mutations, where *i* is either 0 or 1, will be referred to as type-*i* cells or simply, clone *i*. In Materials and Methods we also analyze the more general case of two nested or sibling driver mutations, as well as the fully generalized case of any clonal structure that might arise during tumor expansion.

### Parameter estimates from two longitudinal measurements

We demonstrate that with two longitudinal bulk sequencing measurements, it is possible to accurately estimate net growth rates, time of appearance of a driver mutation, time between a driver mutation and observation, and mutation rate in the tumor. The tumor is first sequenced at time of observation, *t*_1_ + *t*, where both time of driver mutation, *t*_1_, and time from driver mutation to observation, *t*, are yet unknown (Fig. 1b). A second bulk sequencing is performed at *t*_1_ + *t* + *δ*, a known *δ* time units after the tumor is first observed (Fig. 1b). From the bulk sequencing data, the fraction of cells carrying the driver mutation, *α*_1_ and *α*_2_, can be measured at the timepoints *t*_1_ + *t* and *t*_1_ + *t* + *δ*, respectively. We denote total number of cells in the tumor at the two bulk sequencing timepoints as *M*_1_ and *M*_2_. Number of cells in the tumor can be estimated from measurements of tumor volume [45].

Equating expected values of the sizes of type-0 and type-1 population at the two bulk sequencing time points with the measured numbers of cells present in clones 0 and 1, we obtain estimates of the net growth rates of the two subclones:

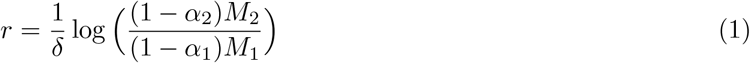

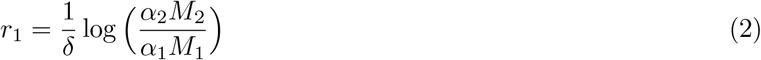

From the growth rate estimates and subclone sizes, we can approximate the expected value of the time a population in a branching process takes to reach an observed size [38]. This yields an estimate of the time *t* from the appearance of driver mutation until observation:

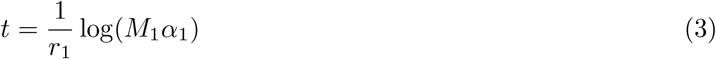

Using the bulk sequencing data from the second timepoint, *γ*, the number of subclonal passengers between the specified frequencies *f*_1_ and *f*_2_, can be measured. Using results from previous work [46], we derive the expected value of *γ* (Materials and Methods), which can be used to estimate the mutation rate *u*:

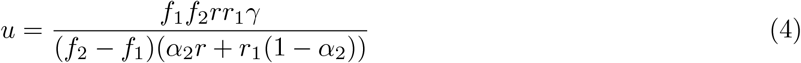

The *m* passenger mutations that were present in the original type-1 cell when the driver mutation occurred (Fig. 1c) are present in all type-1 cells. *m* can be estimated from bulk sequencing data, and used to estimate time of appearance of the driver. We maximize the likelihood function *P* (*m*|*t*_1_) with respect to time of appearance of the driver, *t*_1_, (see Materials and Methods) to obtain the maximum likelihood estimate

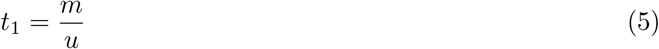

Using formulas (4) and (5), we can now estimate *t*_1_.

### Estimates verified in simulated tumors

To assess the accuracy of the parameter estimates for several modes of tumor evolution, we simulate tumor growth by performing a Monte Carlo simulation, which simulates the birth, death, and accumulation of mutations in the individual cells that make up a tumor. This simulation generates the mutation frequency and tumor size data used by the estimates (see Methods section for details of simulation). We simulate three different types of tumors (slow growing, fast growing, and no cell death), with a high and a low mutation rate for each.

In a simulation of a fast growing tumor with a single subclonal driver mutation that confers a strong selective growth advantage of 100%, we can accurately estimate growth rates, mutation rate, time of driver event, and time since driver event (Fig. 2). Growth rates of both initiated tumor and driver subclones can be estimated with a high degree of accuracy, achieving mean percentage error (MPE) of -0.07% and 0.03% for the lower mutation rate (*u* = 1) scenario. The mutation rate *u* and estimates for time of driver appearance, *t*_1_, and time since driver, *t*, can also be estimated accurately, with MPEs of -0.9%, 3.8%, and -0.4%, respectively. Estimates for *u, t*_1_, and *t* have a somewhat greater degree of variation compared to the growth rate estimates, due to the inherent randomness of the number of mutations and time to reach the observed size that occur in each realization of the stochastic process.

**Figure 2:**
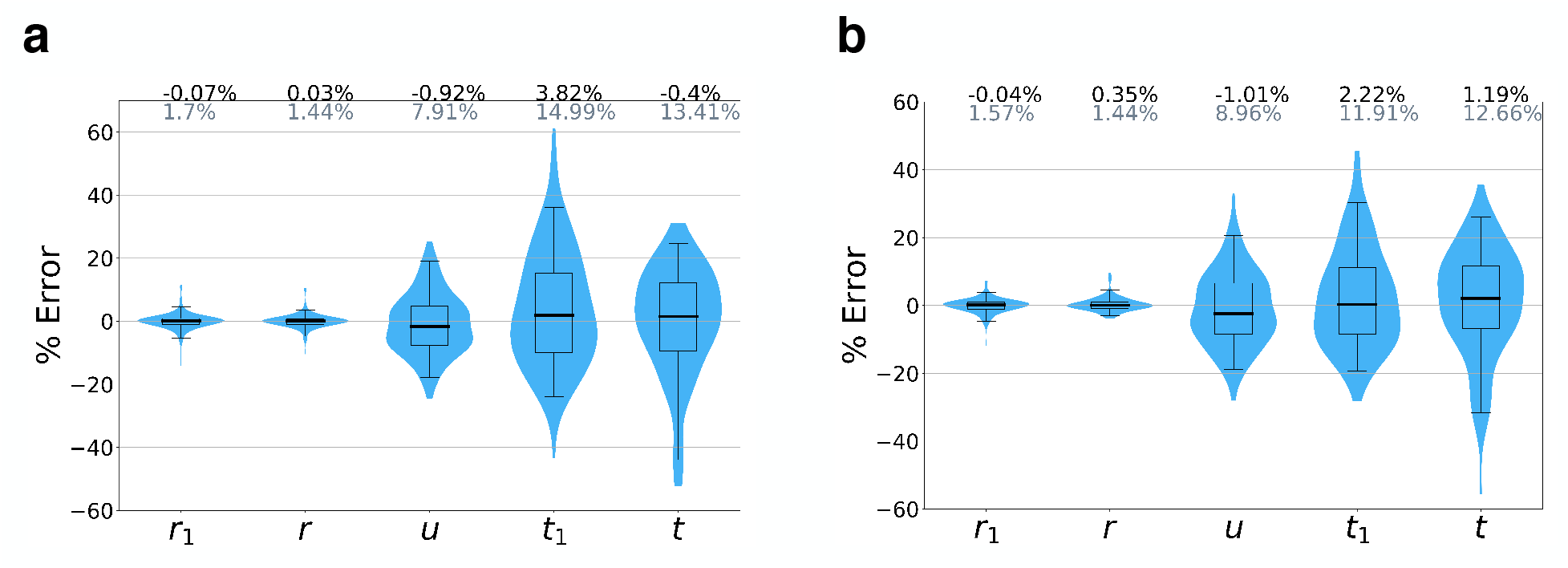
Accuracy of parameter inferences from simulated data. We simulated tumor growth by performing a Monte Carlo simulation, which simulates the birth, death, and accumulation of mutations in the individual cells that make up a tumor, and generates the mutation frequency and tumor size data used by the estimates. Mean percent errors (MPEs) of estimates are shown in black above the plots, and mean absolute percent errors (MAPEs) are shown in gray. Boxes contain 25th-75th quartiles, with median indicated by thick horizontal black line. Whiskers of boxplots indicate 2.5 and 97.5 percentiles. Violins are smoothed density estimates of the percent error data points. Ground truth parameter set: *b* = *b*_1_ = 0.25, *d* = 0.18, *d*_1_ = 0.11, *t*_1_ = 70, *t* = 50, *δ* = 20, *f*_1_ = 1%, and *f*_2_ = 20%. Mutation rate (a) *u* = 1, (b) *u* = 3. At least 100 Monte Carlo simulation runs with a surviving tumor performed for each parameter combination.

For the parameter regime with no cell death and the regime for a slow-growing tumor, we again achieve high accuracies for the net growth rates (Fig. S1, Fig. S2). In the lower mutation rate (*u* = 1) scenario, parameter estimates for the mutation rate *u* and time of driver appearace *t*_1_ can be accurately estimated for both regimes, with MPEs of -1.3% and 4.9% for the no cell death case, and MPEs of -3% and 3.7% for the slow-growing tumor. The *t*, time since driver event, estimates have somewhat higher errors, with MPE of -6.3% for the no cell death case, and MPE of 30.3% for the slow-growing tumor.

### Correcting mutation counts observed from genome sequencing data

We note that in our estimate for the time of appearance of the driver, *t*_1_ (see formula (5)), used for comparison to simulated data, we employed a correction to *m*, the number of mutations that were present in the founder type-1 cell at *t*_1_. From sequencing data, these *m* mutations are indistinguishable (Fig. 3a) from mutations that occurred after *t*_1_ in type-1 cells, and reached fixation in the type-1 population [46]. Thus, the value of *m* observed from sequencing data, *m*_*obs*_, will overestimate the true *m*. In Materials and Methods we show that the expected value of the number of passengers that occurred after *t*_1_ and reached fixation in the type-1 population is *u/r*_1_. We subtract this correction factor from *m*_*obs*_:

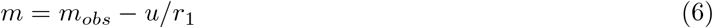

**Figure 3:**
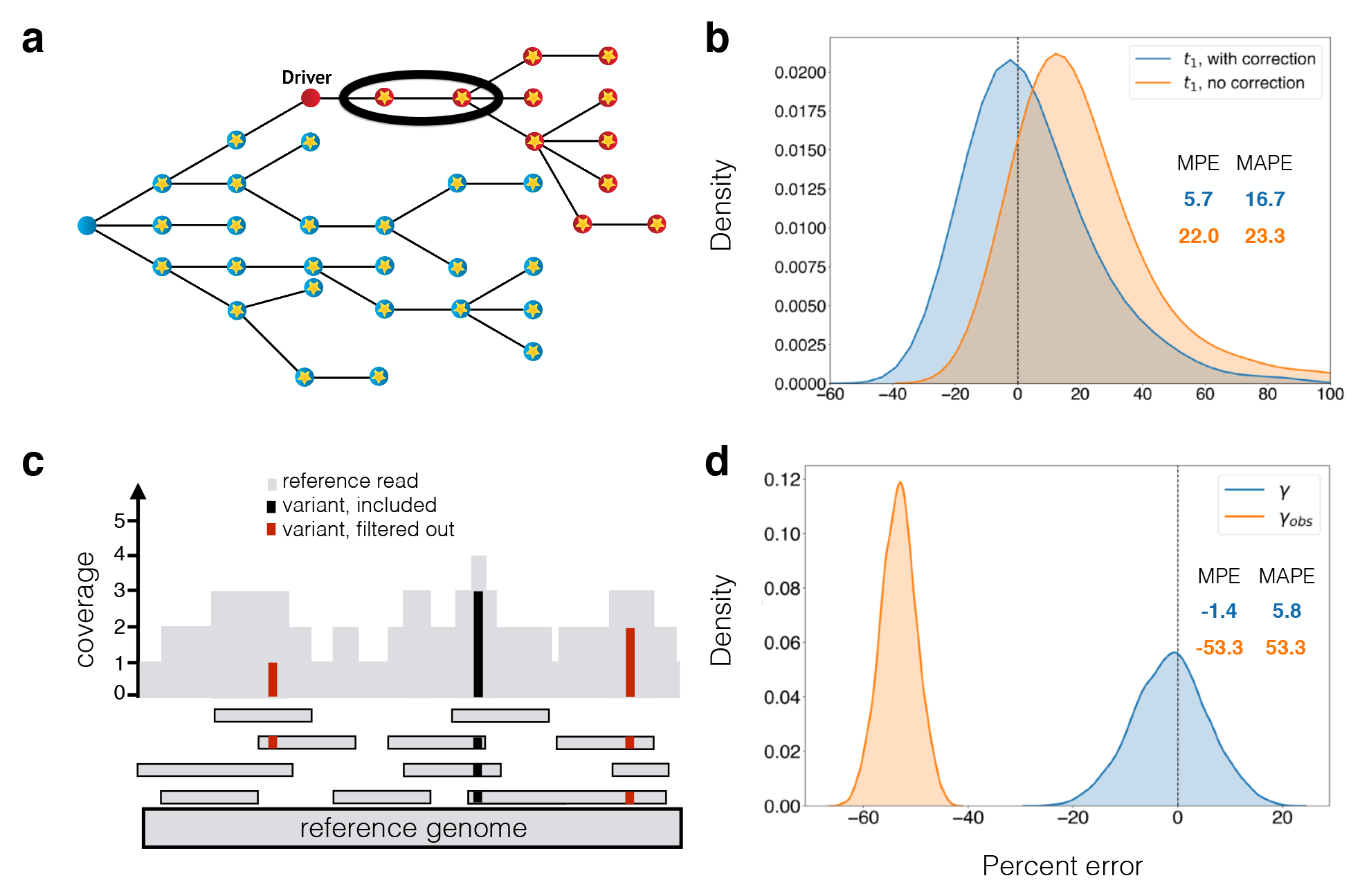
Corrections for observed mutation counts. (a) If passenger mutations (circles with stars) that occur after the driver reach fixation in the driver population (red), then they are indistinguishable from the passengers that were present in the first cell with the driver, which accrued in the type-0 population (blue). The estimate of when the driver occurred needs to account for these mutations (circled). In (b), we compare percent errors of parameter estimates for time from tumor initiaton until appearance of a driver subclone, *t*_1_, with and without this correction (Eq. (6)). Errors for estimate with correction (Eq. (12)) are shown in blue, and for estimate without correction (Eq. (5)) in orange. Errors are plotted as a kernel density estimate for Monte Carlo simulations of slow growing tumor with mutation rate u = 5. Mean percent errors (MPEs) and mean absolute percent errors (MAPEs) are listed. (c) Mutations present on two or fewer variant reads (red) are filtered out in post-processing. Mutations with more than two variant reads (black) are included. The number of subclonal mutations between frequencies *f*_1_ and *f*_2_, *γ*, which is used in the mutation rate estimate, must be corrected for mutations that are filtered out. In (d), the percent errors for the observed (orange) and corrected (blue) *γ* (Eq. (7)) are plotted as kernel density estimates. Observed mutations are those that passed post-processing, i.e. those that have more than *L* = 2 mutant reads. True mutation frequencies were generated from 135 surviving runs of a Monte Carlo simulation of a fast growing tumor with mutation rate *u* = 1, from which sequencing reads were simulated with 200x average coverage (see Materials and Methods). Percent errors are calculated relative to the true *γ* measured from the true mutation frequencies.

The correction for the *m* mutations present in the original type-1 cell (6) at time *t*_1_ improves the accuracy of the estimate for time of appearance of driver mutation *t*_1_. For the fast growing tumor with mutation rate *u* = 1 (Fig. S3a), the correction lowers the mean percent error (MPE) of the *t*_1_ estimate from 14.0% to 3.8%. For the slow growing tumor with mutation rate *u* = 5 (Fig. 3b), the correction lowers the MPE of the *t*_1_ estimate from 22.0% to 5.7% (Fig. 3b).

Another issue arises from obtaining mutation count *γ*, number of mutations with frequency between *f*_1_ and *f*_2_, from genome sequencing data. When sequencing data is post-processed by filtering out mutations with *L* or fewer variant reads, low-frequency mutations will be difficult to detect [35] (Fig. 3c). For a sample with average sequencing coverage of *R* and tumor purity *p*, mutations with mutant allele frequency below *L/*(*pR*) will typically not be observable. As a result, since mutations with frequencies between *f*_1_ and *f*_2_ count towards *γ*, if *f*_1_ ≤ 2*L/*(*pR*), the observed number of subclonal mutations between frequencies *f*_1_ and *f*_2_, *γ*_*obs*_, will underestimate the true value, *γ*. In the Materials and Methods, we derive a correction for *γ*, based on the expected value of the number of subclonal mutations present at cancer cell frequencies (CCFs) between *f*_1_ and 2*L/*(*pR*):

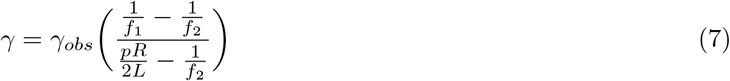

Before applying our methodology to patient sequencing data, we estimated the validity of the above correction applied to observed simulated mutation counts. When we simulate sequencing reads from simulated mutation frequencies (see Materials and Methods) and post-process by removing mutations with *L* = 2 or fewer variant reads, the adjustment we derived for mutation count *γ* (7) is critical, even for average sequencing coverage of 200x (Fig. 3d). Without any correction, the observed *γ* has MPE of -53.3% compared to true *γ*, but with the correction, the computed *γ* has MPE of -1.4%. When average coverage is 100x, this correction becomes even more important, as many of the low-frequency mutations are discarded (Fig. S3b). Without any correction, the observed *γ* has MPE of -79.7%. With the correction the computed *γ* has MPE of -3.4%. The accuracy of the *γ* measurement affects our estimate of the mutation rate (4).

### Estimating parameters for individual patients with CLL

We use our formulas to infer the patient-specific parameters of cancer evolution for four patients with chronic lymphocytic leukemia (CLL) whose growth patterns and clonal dynamics were analyzed in [27]. These CLLs had peripheral white blood cell (WBC) counts measured and whole exome sequencing (WES) performed at least twice before treatment. We consider patients whose WBC counts were classified as having an exponential-like growth pattern, with average *γ*_*obs*_ *>* 2 and 3 or fewer macsoscopic subclones (i.e. subclones with cancer cell fractions of 20% or greater for at least one pre-treatment time point). As in Ref. [27], we perform subclonal reconstruction for each patient using PhylogicNDT [43]. To obtain confidence intervals for our parameter estimates, we utilize a sampling procedure to account for model and measurement uncertainties, including uncertainties in subclone frequencies, fitted growth curves, and the Poisson process for mutation accumulation (see Materials and Methods). For each patient’s tumor, we compute estimates of the growth rate of each clone, exome mutation rate, the times that each subclone arose, and how long each subclone expanded before the tumor was detected (Table S1). We reconstruct these histories for tumors with various clonal structures.

Patients 3 and 21 are examples of a CLL with a single subclone (Fig. 4). For Patient 3, Clone 0, the most recent common ancestor (MRCA) of this patient’s CLL, was initiated when the patient was 14.6 [1.4, 26.8] years old (median and [95% confidence interval] of estimate). Clone 0 grew with a net growth rate of 0.51 [0.20, 0.85] per year. 18.9 years later, Clone 1 was initiated when the patient was 33.5 [24.1, 39.2] years old. Clone 1 expanded with a growth rate of 0.85 [0.65, 1.04] per year (corresponding to a selective growth advantage of 68.7% over Clone 0), and the patient was diagnosed 29.5 [23.8, 38.9] years later at age 63. We find that the CLL exome mutation rate was 0.48 [0.39, 0.59] mutations per year in this patient.

**Figure 4:**
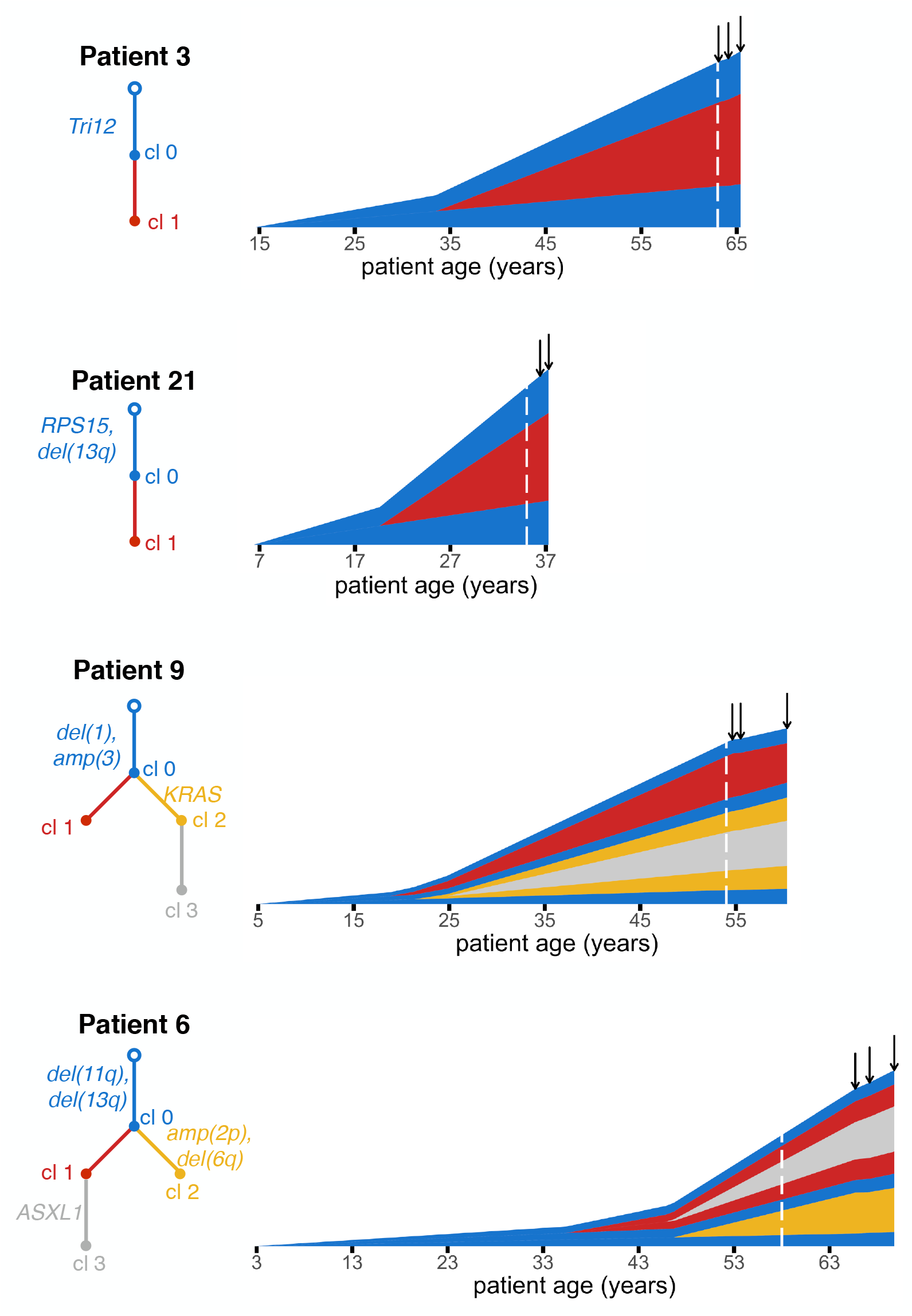
Reconstructing the timeline of CLL evolution in patients. We applied our methodology to estimate subclonal growth rates, mutation rates and evolutionary timelines in CLL tumors from Ref. [27]. Vertical height of a clone represents its log_10_-scaled size. Phylogenetic trees, colored by clone number, show annotated driver mutations along the trees’ edges. For each patient, we show estimates for patient age at CLL initiation and times of appearance of CLL subclones. Dashed white line indicates when the patient was diagnosed. Solid black arrows indicate times of bulk sequencing measurements.

For patient 21, we estimate that the parental clone (MRCA, Clone 0) of this patient’s CLL was initiated when the patient was 6.4 [0.3, 16.7] years old, and grew with a net growth rate of 0.79 [0.30, 1.14] per year. Clone 1 appeared when the patient was 19.6 [10.8, 24.0] years old, and grew more quickly than Clone 0, with a growth rate of 1.52 [1.01, 2.04] per year (corresponding to selective growth advantage of 91.4% over Clone 0). Clone 1 contained a FGFR1 mutation, which might have been acting as a driver of the increased net proliferation. Clone 1 then grew for 15.4 [11.0, 24.2] years before the patient was diagnosed at age 35. We estimate that this patient’s CLL had an exome mutation rate of 0.20 [0.19, 0.23] mutations per year.

Patients 6 and 9 present more complex clonal structures. CLL of Patient 9 contains two sibling subclones, Clones 1 and 2, in addition to the parental population, Clone 0. Clone 2 contains a nested subclone (Clone 3). Clone 0 arose when the patient was 4.9 [1.2, 10.8] years old, and had a growth rate of 0.28 [0.17, 0.42] per year. Clone 1 arose when the patient was 18.8 [8.8, 35.1] years old. At the time of sequencing, Clone 1 had a negative growth rate of -0.40 [-0.45, -0.19] (/year). Clone 2, containing a KRAS mutation, had the largest net growth rate of the three clones (0.67 [0.49, 0.94] per year), corresponding to a selective growth advantage of 140.9% over the parental clone. Clone 2 arose when the patient was 21.3 [7.7, 31.7] years old. Clone 3 was initiated from within Clone 2 when the patient was 24.8 [10.3, 37.6] years old. We estimate that the CLL exome mutation rate of Patient 9 is 0.36 [0.35, 0.37] mutations/year.

CLL of Patient 6 contains two sibling subclones (Clones 1 and 2) descendant from the leukemic MRCA Clone 0. Clone 1 has a nested subclone (Clone 3). We estimate that the CLL was initiated when the patient was 2.8 [0.1, 13.2] years old. Clone 0 then grew at a rate of 0.68 [0.15, 1.30] per year. Approximately 33 years after the appearance of Clone 0, when the patient was 35.4 [21.7, 46.1] years old, the first subclone, Clone 1 appeared. Clone 1 had a net growth rate of 0.41 [0.08, 0.73] per year. Clone 3 arose from within Clone 1 when the patient was 45.9 [31.3, 54.6] years old. This clone had net growth rate 1.09 [0.65, 1.78] per year. Clone 3 harbored a driver mutation in ASXL1 and had selective growth advantage of 60.8% over Clone 0. Clone 2, nested in parental clone (Clone 0), was initiated when the patient was 46.7 [25.6, 57.5] years old and had growth rate 0.46 [0.08, 0.85] per year. The patient was then diagnosed at age 58, eventually needing treatment 12.0 years after diagnosis. In Patient 6, we estimate a CLL exome mutation rate of 0.15 [0.12, 0.19] mutations per year.

The average mutation rate in the four CLL patients we analyze is 0.30 mutations/year. This rate is over the exome, which accounts for ∼ 1% of the human genome. Our average estimated mutation rate in CLL exomes is similar to the measured rate of accumulation of mutations in human tissues of 40 mutations per year over the entire genome [47]. Other recent work has estimated a mutation rate of 17 mutations per year in human haematopoietic stem cell/multipotent progenitors [48]. Our estimated mutation rates during CLL progression are on par or higher than the recent estimates in healthy hematopoietic cells [48], in line with the expectation that mutation rates may be increased in cancer. The estimated times of appearance of CLL subclones are very long, on the order of 10 years or more. This finding is agreement with results from Gruber et al. [27], who find few new CLL subclones over years to a decade of evolution. We observe that CLL initiation occurred early in most patients, within the first fifteen years of their lives, consistent with recent work in other cancer types [19, 36].

## Discussion

We use a stochastic branching process model to reconstruct the timing of driver events and quantify the evolutionary dynamics of different subclonal populations of cancer cells. We can accurately estimate the growth rates of tumor subclones, selective growth advantage of individual driver mutations, mutation rate in the tumor, time between tumor initiation and appearance of a subclonal driver mutation, and time between driver mutation and tumor observation. Together, this allows us to estimate the age of the patient at tumor initiation, as well as the age at appearance of a subclonal driver.

Previous work has computed relative order of driver events [18, 21, 49], while other studies have given estimates for scaled mutation rates and time of events [24, 32]. However, we present estimates for absolute, unscaled mutation rates and times, which are easily interpretable and don’t implicitly depend on unknown parameters. We assume that mutations accrue with time, and not only at cell divisions, which simplifies derivations and is supported by recent experimental data [47].

For individual CLLs that underwent bulk sequencing at two time points [27], we infer growth rates of individual subclones, mutation rate in the tumor, the times when cancer subclones began growing, and the time between driver mutations and the patient’s diagnosis. Our inferences are limited by the relatively low number of mutations present in CLL, as well as sequencing coverage [27]. The accuracy of estimates presented here is expected to be even higher in cancer types with more mutations, with whole genome sequencing available, or with higher sequencing coverage. Our methodology is in principle applicable to any cancer type, not only CLL or liquid cancers. We note, however, that in the case of solid tumors, multiple biopsies would potentially be needed to fully account for the existing heterogeneity.

Our model and derivations assume a fixed mutation rate *u* after transformation and fixed growth rates of cancer subclones, similar to previous approaches [24, 30, 35]. Using an exponential model of cancer growth with constant mutation and growth rate to estimate parameters of cancer evolution has its weaknesses: some cancer subclones (such as Clone 1 from Pt. 9) not only do not grow exponentially, they actually decline in absolute cell numbers. Sudden genomic instability events, or a change in cancer mutation and/or growth rate over time could also introduce errors into our parameter inferences. Recent sequencing data points to mutational processes that change over time during cancer evolution [20, 50]; incorporating possible changes in the mutation and/or growth rate into the model would require much higher density of sequencing and clinical data [37].

## Materials and Methods

### Branching process model of tumor evolution

We employ a continuous, multi-type branching process model of cancer evolution. Tumor expansion is initiated by a single type-0, or initiated tumor cell. Type-0 cells divide with rate *b* and die with rate *d*, yielding a net growth rate of *r* = *b* − *d*. At time *t*_1_, a single driver mutation is introduced into a randomly selected cell in the type-0 population, founding a new type-1 population of cells. This type-1 population undergoes its own independent branching process. They divide with rate *b*_1_, die with rate *d*_1_, and have net growth rate *r*_1_ = *b*_1_ − *d*_1_. If the driver mutation gives type-1 cells a selective growth advantage over the type-0 population, then *r*_1_ *> r*. With the ratios of the growth rates denoted as *s* = *r*_1_*/r*, the growth advantage can be quantified as *g* = (*s*−1)·100%. In the case of neutral evolution, *g* = 0. If there is a selective advantage, *g >* 0. Neutral mutations, or passengers, have no effect on the cell’s fitness, and accrue according to a Poisson process with rate *u*. We assume an infinite alleles model such that there is no back mutation and an infinite sites model such that every new passenger mutation is unique. Only surviving populations are considered. All derivations below will condition on survival. The type-0 and type-1 populations at time *t* will be denoted as *X*_0_(*t*) and *X*_1_(*t*), respectively.

### Measurements sufficient to determine evolutionary history

We derive estimates for parameters describing the carcinogenic process using measurements taken from two timepoints late in the tumor’s development. We require sequencing of the tumor at the two timepoints, when the tumor is first observed at the unknown time *t*_1_ + *t* and a specified *δ* later, at *t*_1_ + *t* + *δ*. From these two bulk sequencing measurements, we obtain measurements of *α*_1_ and *α*_2_, the fraction of cells carrying the driver mutation at *t*_1_ + *t* and *t*_1_ + *t* + *δ*, respectively. In addition, from the bulk sequencing at *t*_1_ + *t* + *δ*, we obtain measurements of *m*, the number of mutations present in the founder type-1 cell, as well as *γ*, the number of mutations with frequency between the specified *f*_1_ and *f*_2_. The total population size at these times, *M*_1_ and *M*_2_, is also measured.

### Expected value of *γ*, number subclonal mutations

For a population consisting of a single clone with birth and death rates *b* and *d*, the expected number of subclonal mutations present at a frequency larger than *f* is shown to be [46]

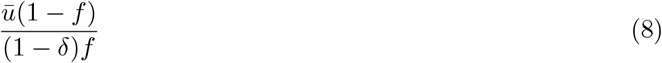

where *δ* = *d/b* and *ū* is the probability that a daughter cell gains a new passenger mutation at cell division. In this paper, we allow mutations to occur at any point in time and consider the absolute mutation rate per cell, *u*, which is equal to *ūb*. Then the expected number of subclonal mutations between *f*_1_ and *f*_2_, 𝔼*γ*, is

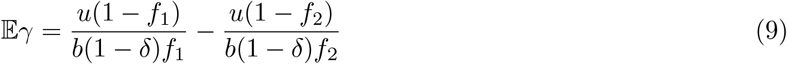

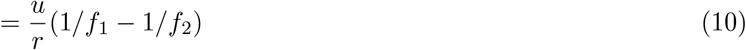

where *r* = *b* − *d >* 0.

Now we derive 𝔼*γ* in the case of clones 0 through *k*, each clone with growth rate *r*_*i*_ *>* 0 and fraction 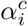. Each clone *i* has 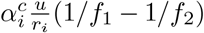 expected subclonal passengers between frequencies *f*_1_ and *f*_2_. Thus, the total expected number of passengers with frequencies between *f*_1_ and *f*_2_ is

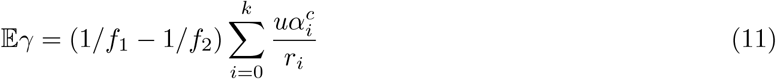

For the simplest case we consider, a tumor with a single driver mutation occurring in the initiated tumor population, there is a type-0 population with growth rate *r* and a type-1 population with growth rate *r*_1_. Equation (11) reduces to

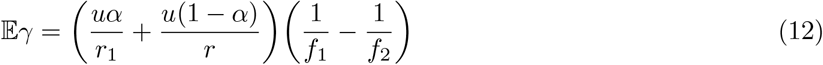

where *α* is the fraction of cells having the driver mutation.

### Derivation of estimates of evolutionary parameters

With the two bulk sequencings at *t*_1_ + *t* and *t*_1_ + *t* + *δ*, we are able to derive estimates for *t*_1_, *t, r, r*_1_, and *u*. First we solve for *r* and *r*_1_, based on the estimated cell counts at *t*_1_ + *t* and *t*_1_ + *t* + *δ*. The observed type-*i* cell count is equated to the expected value of the type-*i* population size, conditioned on survival. For the type-0 population,

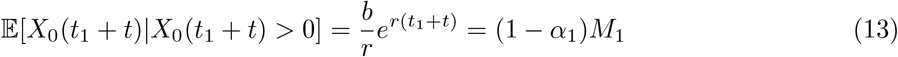

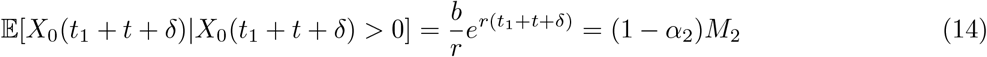

Proceeding similarly for the type-1 population, we obtain

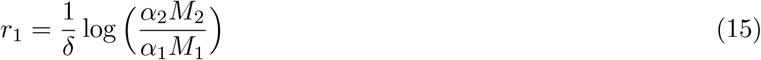

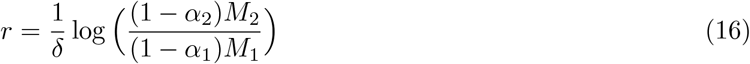

The expected value of the first time a population of type-1 cells in a branching process reaches the observed size *α*_1_*M*_1_ is [38]

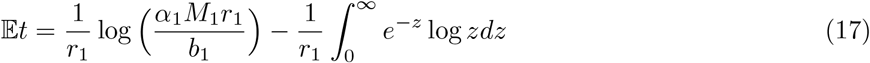

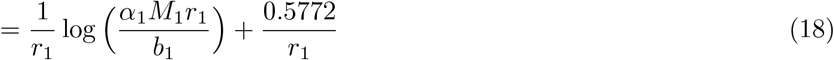

which we approximate as

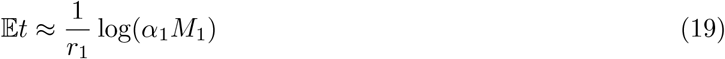

We make use of two approximations to arrive at (19). First, we neglect the second term in (18), which serves as a small correction to the first term. This term will be dominated by the first term as it increases with logarithm of the cancer size. For *r*_1_ = 0.5, *α*_1_*M*_1_ ∼ 10^11^, and *r*_1_ ≈ *b*_1_, the second term (1.2) will be only 2.3% of the first term (50.7). For any growth rate, the second term will be 2.3% of the first term. Second, we assume *r*_1_ is similar in magnitude to *b*_1_.

With the measurement of *γ*, the number of subclonal passengers with frequency between *f*_1_ and *f*_2_, we can estimate the mutation rate *u*. In the previous section we derive the expected value of *γ* as

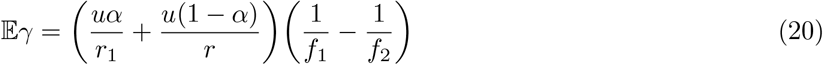

Using the estimates of *r* and *r*_1_ from (15) and (16), and the measured value of *γ* from the second bulk sequencing, equation (20) can be solved for the mutation rate *u*,

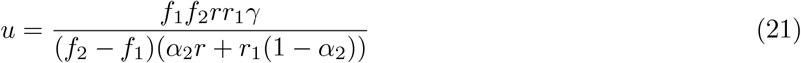

When estimating mutation rate for the CLL patients from Ref. [27], for which there is bulk sequencing at two or more timepoints, we average the mutation rate calculated at each of these timepoints. (21) is applied for each timepoint with the respective CCFs and observed *γ* values for each timepoint.

To derive the maximum likelihood estimates of *t*_1_, we consider the likelihood function *P* (*m*|*t*_1_). The number of passenger mutations present in the founder type-1 cell that appeared at time *t*_1_ is a Poisson process with rate *u*. Thus,

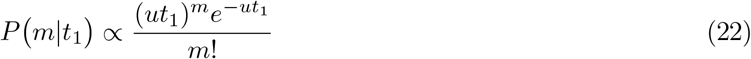

Maximizing the logarithm of the likelihood function with respect to *t*_1_ yields a MLE for *t*_1_ in terms of estimated or measured quantities:

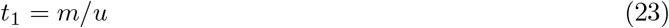

### Estimating number of unobserved subclonal mutations from sequencing data

When sequencing data is post-processed by filtering out any mutations with *L* or fewer variant reads, the number of mutations between *f*_1_ and *f*_2_ will likely be underestimated if 2*L/*(*Rp*) *> f*_1_, where *R* is average sequencing coverage and *p* is tumor purity. Define *γ*_*obs*_ as the observed number of mutations between frequencies *f*_1_ and *f*_2_, after post-processing has been performed that filtered out any mutations with *L* or fewer variant reads. The expected number of subclonal mutations between frequencies *f*_1_ and *x* is given by

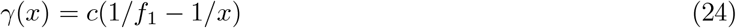

where *c* is a constant that will vary depending on the patient and sample. It can be fit on the sequencing data by noting

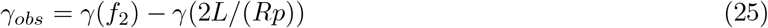

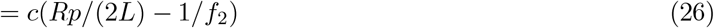

Therefore, *c* can estimated from the sequencing data as

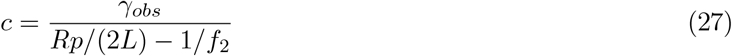

Then, we can estimate *γ* as

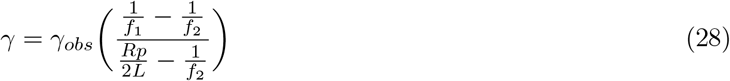

### Number of passengers reaching fixation after *t*_1_

We estimate the number of passengers that occurred after *t*_1_ and reached fixation in the type-1 population in order to adjust the *m*_*obs*_ mutation count. From [46], when mutations occur at cell division, the expected number of clonal passengers is *δū/*(1 − *δ*). *ū* is the probability that a daughter cell gains a new passenger mutation at cell division, so the mutation rate is *u* = *ūb*_1_. For the type-1 population, *δ* = *d*_1_*/b*_1_ *<* 1. When mutations accrue over time, and not only at divisions, the expected number of clonal passengers is thus

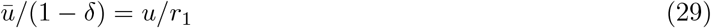

Similarly, for a clone *i*, the expected number of passengers that occur after time *t*_*i*_ and reach fixation is

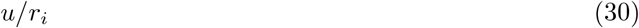

where *r*_*i*_ = *b*_*i*_ − *d*_*i*_ *>* 0.

### Simulation of tumor evolution and sequencing data

To assess the accuracy of the analytic results, we perform a continuous time Monte Carlo simulation to model tumor evolution and collection of sequencing data with an implementation of the Gillespie algorithm [51]. Simulations are written in C/C++.

The type-*j* population has division rate *b*_*j*_, death rate *d*_*j*_, and mutation rate *u*. Mutations can occur at any point of the cell cycle, not just during division. *z*_*n*_ is the number of type-*j* cells with passenger *n* as their most recent passenger mutation. The type-0 population is initiated with a single cell at time 0, and the type-1 population is initiated with a single cell at time *t*_1_. Let *a* be the vector recording the ancestor of new mutations. Element *a*_*i*_ is the subclonal ancestor of the *i*th passenger mutation. Repeat 1-4 while time is less than *t*_1_ + *t* + *δ*.

1. Set Γ = *N*_*j*_(*b*_*j*_ + *d*_*j*_ + *u*). Time increment to next event time is randomly sampled from Exp[Γ].
  - For type-0, if time is greater than or equal to *t*_1_ for first time, randomly select type-0 subclone *i* to have driver mutation, remove one cell from type-0 population count, and set *N*_1_ = 1. Record the true value of *m*, the number of passenger mutations present in the founder type-1 cell.
2. Randomly select cell, with most recent passenger mutation *i*, to have the event.
3. Determine which type of event and update population and mutation frequencies. Sample *Y* from Uniform[0, Γ] to determine event type:
  i. *y* ∈ (0, *b*_*j*_) → birth. *N*_*j*_ += 1, *z*_*i*_ += 1.
  ii. *y* ∈ (*b*_*j*_, *b*_*j*_ + *d*_*j*_) → death. *N*_*j*_ −= 1, *z*_*i*_ −= 1.
  iii. *y* ∈ (*b*_*j*_ + *d*_*j*_, *b*_*j*_ + *d*_*j*_ + *u*) → passenger mutation. Suppose it’s the *k*th passenger, *z*_*i*_ −= 1, *z*_*k*_ = 1. Update ancestor: *a*_*k*_ = *i*.
4. For type-0, if time is less than *t*_1_ and population goes extinct, restart simulation. For type-1, if time is greater than *t*_1_ and population goes extinct, restart type-1 simulation at *t*_1_ with a single cell.
5. Reindex to remove extinct passenger mutations, and traverse back through ancestor vector **a** to sum total number of cells with each passenger.

Measurements are taken at bulk sequencing times *t*_1_ + *t* and *t*_1_ + *t* + *δ*. If time is greater than or equal to *t*_1_ + *t*, we measure *M*_1_ = *N*_0_ + *N*_1_ and *α*_1_ = *N*_1_*/*(*N*_0_ + *N*_1_). Then an additional bulk sequencing measurement is taken at the final time *t*_1_ + *t* + *δ*, where we measure *M*_2_ = *N*_0_ + *N*_1_ and *α*_2_ = *N*_1_*/*(*N*_0_ + *N*_1_). At *t*_1_ + *t* + *δ*, we measure *γ*, the number of mutations with frequency between *f*_1_ and *f*_2_.

To measure *m*_*obs*_, the observed number of passengers in the founder type-1 cell, we count the number of passengers present in all type-1 cells. We also save the true value of *m*.

For when we calculate a percent error of corrected and observed *γ* values in Figure 3d and Supplementary Figure 3b, we simulate sequencing data by sampling from the mutation frequencies obtained in the Monte Carlo simulation, outlined above, using the approach of [35]. Define average sequencing coverage as *R*, number of cells at time of sequencing as *M, Z*_*i*_ as the number of cells with mutation *i, R*_*i*_ as read coverage, and *χ*_*i*_ as the true mutation frequency from Monte Carlo simulation. For each saved Monte Carlo simulation run, repeat the following 100 times:

1. Generate read coverage: *R*_*i*_ ∼ Binomial[*M, R/M* ]
2. Generate number of cells carrying mutation *i*: *Z*_*i*_ ∼ Binomial[*R*_*i*_, *χ*_*i*_*/*2]
3. Post-processing. If there are *L* = 2 or fewer variant reads, discard mutation.
4. Measure *γ*_*obs*_, the observed number of subclonal mutations between frequencies *f*_1_ and *f*_2_: *γ*_*obs*_ = Σ_*i*_ *I*(*f*_1_ ≤ 2*Z*_*i*_*/R* ≤ *f*_2_, *Z*_*i*_ *> L*)
5. Calculate the truth, *γ*_*true*_, from the true mutation frequencies: *γ*_*true*_ = Σ_*i*_ *I*(*f*_1_ ≤ *χ*_*i*_ ≤ *f*_2_)

### Parameter values for simulations

For the simulation we consider three parameter sets corresponding to three modes of tumor evolution: a fast growing tumor, slow growing tumor, and tumor with no cell death. For each parameter regime we have a low and high mutation rate. Mutation rate parameter values lie within observed genome wide point mutation rates per day [52]. For the fast growing tumor *b* = *b*_1_ = 0.25, *d* = 0.18, *d*_1_ = 0.11, *t*_1_ = 70, *t* = 50, *δ* = 20, and *u* = 1, 3. For the slow growing tumor *b* = 0.25, *b*_1_ = 0.25, *d* = 0.225, *d*_1_ = 0.2125, *t*_1_ = 180, *t* = 135, *δ* = 45, and *u* = 1, 5. For the parameter regime with no cell death *b* = 0.25, *b*_1_ = 0.375, *d* = *d*_1_ = 0.0, *t*_1_ = 23, *t* = 17, *δ* = 6, and *u* = 1, 10. The fast growing tumor dynamics are from [34]. The slower growing tumor parameter regime has a reduced net growth of *r* = 0.025, compared to the fast growing tumor’s net growth rate of *r* = 0.07.

### Subclonal reconstruction of CLL sequencing data

The sequencing data from all CLLs analyzed is from Ref. [27], Supplementary Tables 2-4. As in that publication, we use PhylogicNDT [43] to perform subclonal reconstruction. We run the Cluster and BuildTree modules of PhylogicNDT on the longitudinal mutation data from Supplementary Table 3 of [27], using mutation alternate/reference counts, copy number, and tumor purity at all pre-treatment time points. Then for each patient, PhylogicNDT outputs a clonal reconstruction, which includes a phylogenetic tree of the subclones and posterior distributions of subclone CCFs. Additionally, it clusters mutations and assigns them to clones. We directly use subclone assignments and posteriors generated from PhylogicNDT. In our analysis we focus on estimating timing and growth rates of macroscopic subclones whose CCFs are greater than 20% for at least one pre-treatment timepoint.

### Accounting for uncertainties in subclone frequencies and growth rates

Our estimates for parameters of cancer evolution require as input the information on the number of subclonal populations in the tumor, their CCFs and their phylogenetic relationships. In order to obtain this information, we use PhylogicNDT [43], which performs subclonal reconstruction of longitudinal cancer sequencing data. The uncertainty in subclone CCFs reported by PhylogicNDT affects our estimates for subclone growth rates, which in turn affect the estimates of mutation rate and and time *t* between driver(s) and diagnosis. We account for this uncertainty by drawing from the CCF posterior distributions that are output by PhylogicNDT. Using these sampled CCF values, we then calculate growth rates, mutation rate *u*, and time *t* between driver(s) and diagnosis, thereby generating confidence intervals for these parameters due to CCF uncertainty.

To estimate subclonal growth rates, we fit an exponential growth curve to subclonal sizes measured at two or more time points. This regression yields fitted values for each clone’s growth rate and age. To account for uncertainty in the curve fit (in the case of more than two longitudinal samples), we sample the growth rates and age of clone from a bivariate normal distribution with mean equal to the fitted parameters and variance equal to the covariance matrix of the fitted parameters. In line with recent findings [53], we found that sometimes the estimated growth rate is smaller than minimal possible growth rate necessary to reach the observed clone size. In that case, for calculating mutation rate, time of the driver(s), and time between driver(s) and diagnosis we use the minimal growth rate.

### Accounting for model uncertainty

The largest source of model uncertainty is the Poisson process for how mutations accumulate, which is used to estimate the time *t*_1_ of the driver mutation. In the simulation experiments, the time *t*_1_ had the largest error and variation (Fig. 2). The estimate for *t*_1_ depends on the *m* mutations present in all cells in the driver subclone. The observed *m* is a single random sample from a Poisson distribution. To account for the uncertainty in *t*_1_ arising from *m* in the CLLs analyzed, we sample *t*_*i*_ from the posterior distributions *P* (*t*_1_|*m*). This source of model uncertainty due to the Poisson process will be most significant for cancers like CLL with a smaller number of mutations.

The time *t* between driver mutation and diagnosis (*t*) is a random variable due to the stochasticity of cancer cell growth, and will naturally have a certain amount of variation. Time between driver event and diagnosis in a branching process follows a Gumbel distribution [38], and will have a constant variance. The mean, however, will increase with the logarithm of the cancer cell counts, which for the CLLs analyzed are ∼ 10^11^. The simulations of cancer evolution grow to smaller tumor sizes (∼ 10^5^) and, as a result, the estimate for *t* has a significant amount of uncertainty (Fig. 2). However, for time scales necessary to generate a tumor, the estimate for *t* will be quite accurate. For commonly observed tumor sizes, the stochastic fluctuations in the time for the cancer to reach that size will be smaller relative to the magnitude of the time. For a cancer with cell count ∼ 10^11^, the standard deviation of the time *t* will be less than 5% of its expected value.

### Tumor with two nested driver subclones

Here we consider the case where there are two nested driver subclones (Fig. S4a). “Nested” means that all cells carrying the second driver mutation also carry the first. Type-0, or initiated tumor, cells have birth rate *b*_0_, death rate *d*_0_, and net growth rate *r*_0_ = *b*_0_ − *d*_0_. Type 1 cells, which only have the first driver, have birth rate *b*_1_, death rate *d*_1_, and net growth rate *r*_1_ = *b*_1_ − *d*_1_. Type-2 cells, which carry both drivers, have birth rate *b*_2_, death rate *d*_2_, and net growth rate *r*_2_ = *b*_2_ − *d*_2_. The first driver occurred in a type-0 cell at time *t*_1_. The second driver occurred in a type-1 cell at 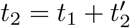. The mutation rate *u* is the same for all subclones.

At times 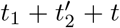 and 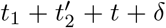, the tumor is bulk sequenced. The bulk sequencing allows the measurement of the fraction of cells with driver 1 at time 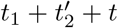, *α*_1_; the fraction of cells with driver 2 at 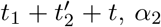; fraction of cells with driver 1 at time 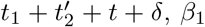; the fraction of cells with driver 2 at 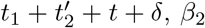; and the observed number of subclonal passenger mutations between frequencies *f*_1_ and *f*_2_, *γ*_*obs*_. Note that the fraction of the population that is a type-1 cell at the two times is *α*_1_ − *α*_2_ and *β*_1_ − *β*_2_. The fraction of type-0 cells at the two bulk sequencing timepoints are 1 − *α*_1_ and 1 − *β*_1_. The number of total cells at bulk sequencing timepoints are *M*_1_ and *M*_2_. Equating the estimated cell counts to the expected value of the type-*i* population size *X*_*i*_, conditioned on survival,

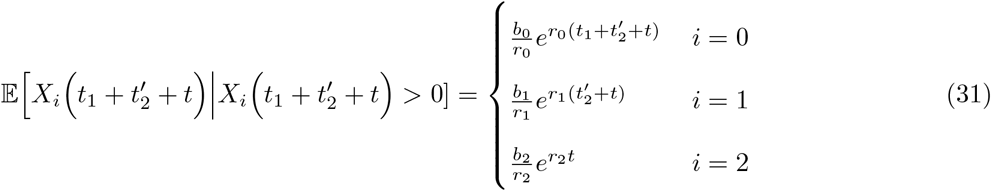

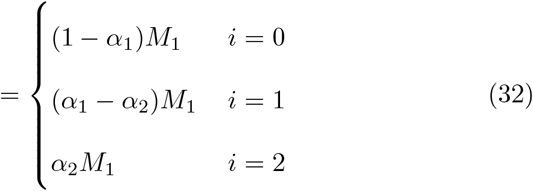

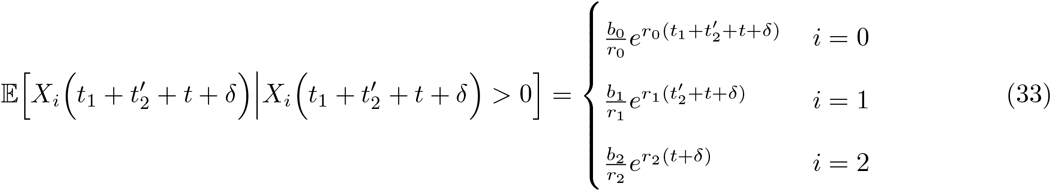

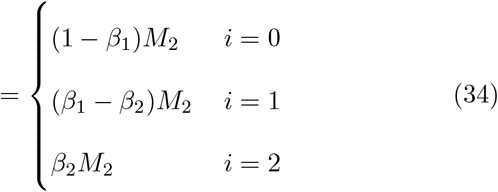

Solving the above equations for *r*_*i*_, we obtain the growth rate estimates:

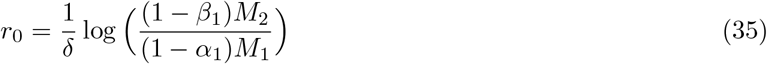

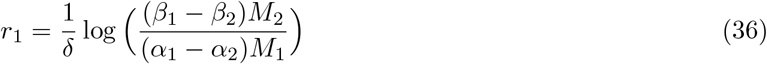

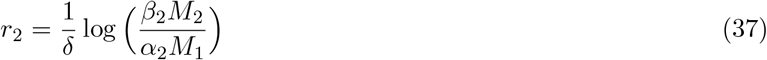

The expected value of the first time a population of type-2 cells in a branching process reaches the observed size *α*_2_*M*_1_ [38],

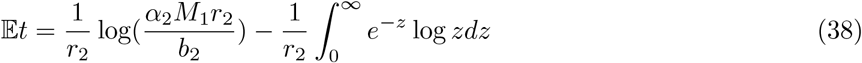

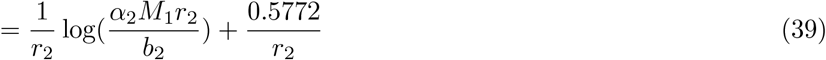

can be approximated as

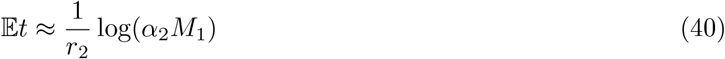

We make use of two approximations to arrive at (40). First, we neglect the second term in (39), which serves as a small correction to the first term. Second, we assume *r*_2_ is similar in magnitude to *b*_2_.

By (11),

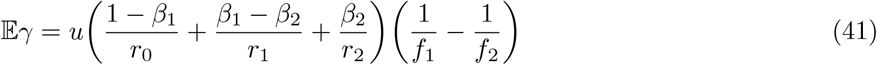

Using the estimates for *r*_0_, *r*_1_, and *r*_2_ from (35)-(37), and setting (41) equal to the value of *γ* obtained from (28) and the second bulk sequencing, *u* can be estimated:

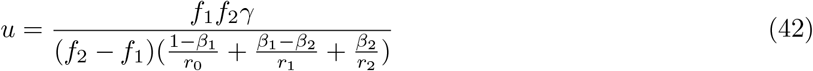

When estimating mutation rate for the CLL patients from Ref. [27], for which there is bulk sequencing at two or more timepoints, we average the mutation rate calculated at each of these timepoints. (42) is applied for each timepoint with the respective CCFs and observed *γ* values for each timepoint.

Every type-1 cell carries the *m*_1_ passenger mutations that were present in the original type-1 cell when the first driver mutation mutation occurred at *t*_1_. Similarly, every type-2 cell carries the *m*_2_ passengers that were present in the founder type-2 cell when the second driver mutation occurred at *t*_2_. Note, none of the *m*_1_ mutations are counted towards *m*_2_. Now we consider the likelihood function

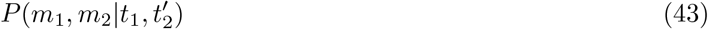

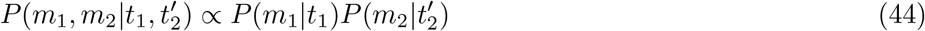

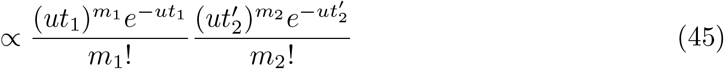

Now, maximizing the logarithm of (45) with respect to *t*_1_ and 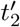,

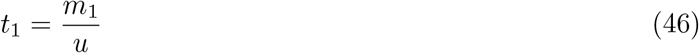

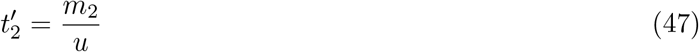

The number of passengers present in the founder type-*i* cell cannot be directly observed, but we can measure *m*_*i obs*_, the number of passengers present in all type-*i* cells. An expected *u/r*_1_ passengers occurring after *t*_1_ in type-1 cells and reaching fixation in the type-1 subclone will be incorrectly included in *m*_1 *obs*_, rather than in *m*_2 *obs*_ (see Methods). Similarly, an expected *u/r*_2_ passengers occurring after *t*_2_ in type-2 cells and reaching fixation in the type-2 subclone will be incorrectly included in *m*_2 *obs*_. Thus,

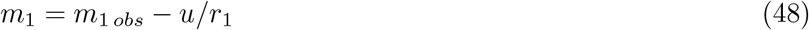

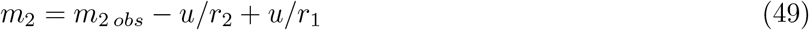

### Tumor with two sibling driver subclones

Here we consider a tumor with two “sibling” driver mutations (Fig. S4b). Sibling driver mutations are drivers that occur in separate subclones. In this case, cells are either initiated tumor cell (type-0), carry driver 1 (type-1), or carry driver 2 (type-2). No cells contain both drivers. Driver 1 occurred in a type-0 cell at time *t*_1_. Driver 2 occurred in a type-0 cell at *t*_2_. Type-0 cells have birth rate *b*_0_, death rate *d*_0_, and net growth rate *r*_0_ = *b*_0_ − *d*_0_. Type-1 cells, which carry driver 1, have birth rate *b*_1_, death rate *d*_1_, and net growth rate *r*_1_ = *b*_1_ − *d*_1_. Type-2 cells, which carry driver 2, have birth rate *b*_2_, death rate *d*_2_, and net growth rate *r*_2_ = *b*_2_ − *d*_2_. The mutation rate *u* is the same for all subclones.

Suppose time *τ*_*i*_ elapses between driver mutation *i* and tumor observation. Bulk sequencing of the tumor is performed at *t*_1_ + *τ*_1_ (or equivalently *t*_2_ + *τ*_2_), and a known *δ* later. Sequencing the tumor allows the measurement of the fraction of cells with driver 1 at the first sequencing, *α*_1_; the fraction of cells with driver 2 at the first sequencing, *α*_2_; fraction of cells with driver 1 at the second sequencing, *β*_1_; the fraction of cells with driver 2 at the second sequencing, *β*_2_; and the number of subclonal passenger mutations between frequencies *f*_1_ and *f*_2_, *γ*. The fraction of type-0 cells at the two bulk sequencing timepoints are 1 − *α*_1_ − *α*_2_ and 1 − *β*_1_ − *β*_2_. The number of total cells at the two sequencing timepoints are *M*_1_ and *M*_2_.

Equating the estimated cell counts to the expected value of the type-*i* population size *X*_*i*_, conditioned on survival,

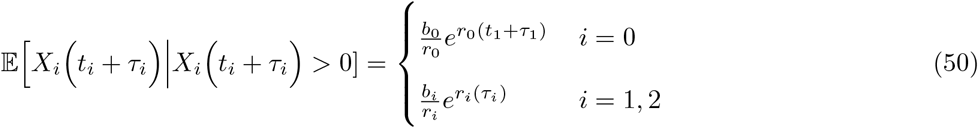

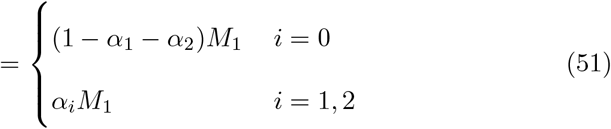

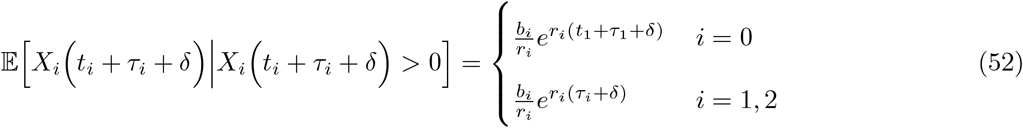

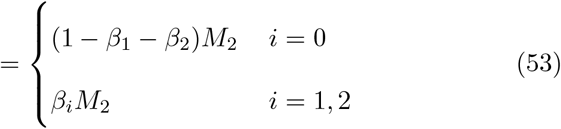

Solving the above equations for *r*_*i*_, we obtain

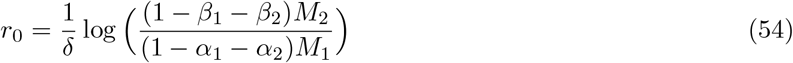

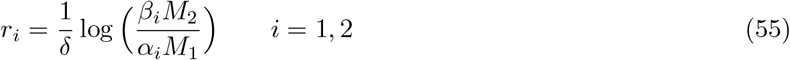

The expected value of the first time a population of type-*i* cells in a branching process reaches the observed size *α*_*i*_*M*_1_ is [38]

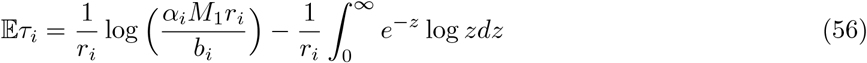

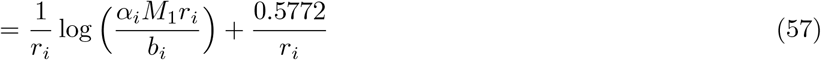

which we approximate as

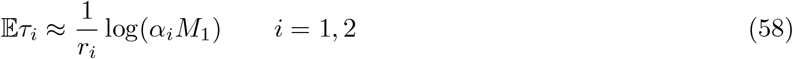

We use two approximations to arrive at (58). We neglect the second term in (57), which serves as a small correction to the first term. Second, we assume *r*_*i*_ is similar in magnitude to *b*_*i*_.

By (11),

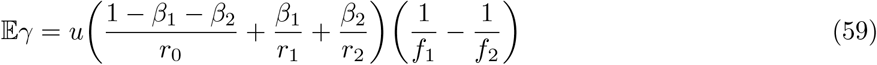

Using the estimates for *r*_0_, *r*_1_, and *r*_2_ from (54) and (55), and setting (59) equal to the value of *γ* obtained from (28) and the second bulk sequencing, *u* can be estimated:

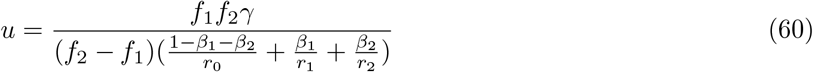

When estimating mutation rate for the CLL patients from Ref. [27], for which there is bulk sequencing at two or more timepoints, we average the mutation rate calculated at each of these timepoints. (60) is applied for each timepoint with the respective CCFs and observed *γ* values for each timepoint.

Every type-1 cell carries the *m*_1_ passenger mutations that were present in the original type-1 cell when the first driver mutation mutation occurred at *t*_1_. Similarly, every type-2 cell carries the *m*_2_ passengers that were present in the founder type-2 cell when the second driver mutation occurred at *t*_2_. We assume that *m*_1_ and *m*_2_ don’t contain any shared mutations. In the CLL dataset we use, this is true. We consider the likelihood function *P* (*m*_1_, *m*_2_|*t*_1_, *t*_2_)

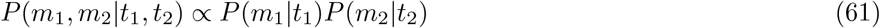

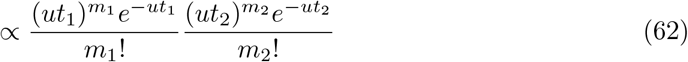

Maximizing the logarithm of (62) with respect to *t*_1_ and *t*_2_ yields the maximum likelihood estimates:

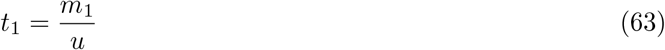

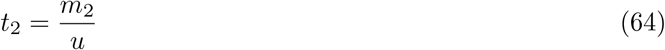

Using the same approach as in the case of a single driver, we obtain the corrections for the observed number of mutations present in all cells of each subclone:

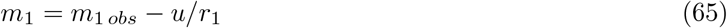

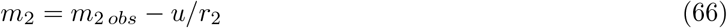

### Fully generalized estimates for any phylogeny of *k* drivers

Here we derive estimates for a completely general tumor phylogeny. Suppose a tumor has *k* driver mutations. In this general case, define a type-*i* cell as a cell where its most recent driver mutation was driver *i*. Note that a type-*i* cell can have between 0 and *k* − 1 other driver mutations. A phylogenetic reconstruction of the *k* driver mutations is necessary for the completely general case. From this phylogenetic tree, the ancestor of each subclone can be obtained. Define the function *a*(*i*) as the ancestor of the type-*i* population. That is, if all driver mutations contained in the type-*i* population are ordered, *a*(*i*) gives the driver mutation that occurred prior to *i*. Define *t*_*i*_ as the time between when driver *i* occurred and when the type-*i* cells’ previous driver mutation occurred. At time of observation, assume the type-*i* population has *κ*_*i*_ total driver mutations, where 1 ≤ *κ*_*i*_ ≤ *k* for all 1 ≤ *i* ≤ *k*. Denote the time between the type-*i*’s *κ*_*i*_, or last, driver mutation and when the tumor is observed as *τ*_*i*_. This is the time between the founder type-*i* cell’s birth and tumor observation. Then the tumor is first observed and bulk sequenced at 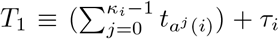 (equivalently *τ*_0_ for *i* = 0), where we denote *a*^*j*^ as the *j*th iterate of the function *a*:

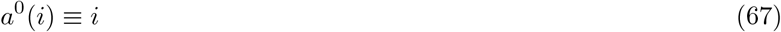

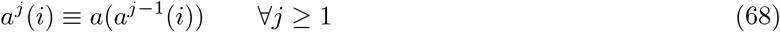

The tumor is also bulk sequenced at 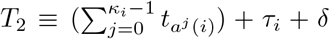 (equivalently *τ*_0_ + *δ* for *i* = 0). These assumptions allow for any subclone phylogeny, including combinations of the previously discussed sibling and nested subclone types.

The bulk sequencing allows the measurement of the fraction of cells with driver *i* at *T*_1_, *α*_*i*_; the fraction of cells with driver *i* at time *T*_2_, *β*_*i*_; and the number of subclonal passenger mutations between frequencies *f*_1_ and *f*_2_, *γ*. Again, the number of total cells at measurement times *T*_1_ and *T*_2_ are *M*_1_ and *M*_2_. To write the type-*i* frequencies, 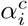 and 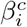, in terms of the driver frequencies, we subtract the fraction of cells descending from type-*i* cells but gaining additional driver mutation(s) after *i*, from the fraction of cells containing driver *i*:

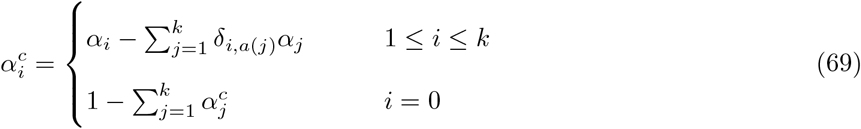

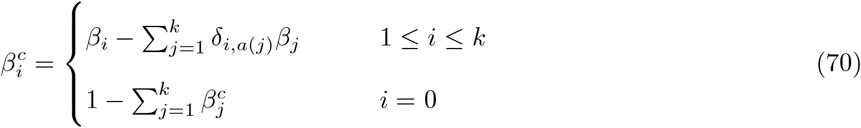

where *δ*_*i,a*(*j*)_ is the Kronecker delta, defined as

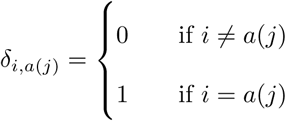

Equating the estimated cell counts at the first bulk sequencing timepoint to the expected value of the type-*i* population size *X*_*i*_, conditioned on survival,

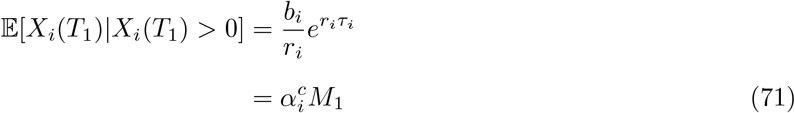

And similarly, at the second bulk sequencing timepoint,

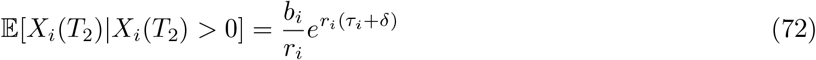

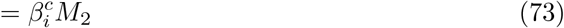

Solving the above equations for *r*_*i*_, we obtain

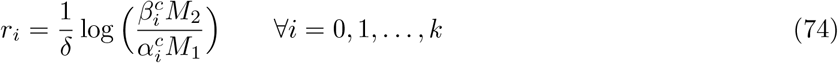

By (11)

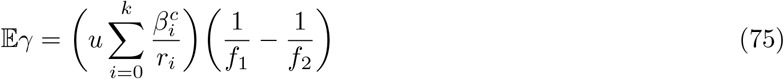

Now, using the growth rate estimates *r*_*i*_ and the subclone sizes, we can estimate each *τ*_*i*_. The expected value of the first time a population of type-*i* cells in a branching process reaches the observed size 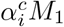 is [38]

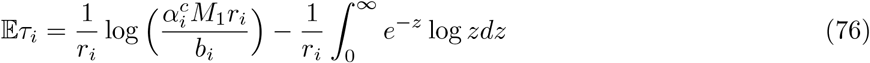

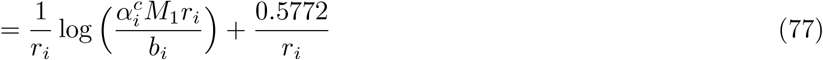

which we approximate as

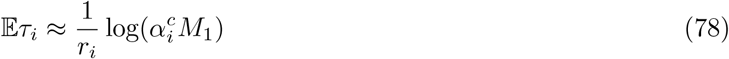

We make use of two approximations to arrive at (78). First, we neglect the second term in (77), which serves as a small correction to the first term. Second, we assume *r*_*i*_ is similar in magnitude to *b*_*i*_.

Using the (*k* + 1) *r*_*i*_ estimates from (74), and setting (75) equal to the value of *γ* obtained at the second bulk sequencing from (28), *u* can be estimated:

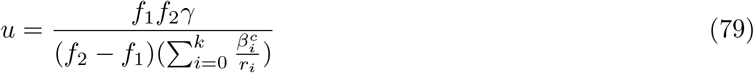

When estimating mutation rate for the CLL patients from Ref. [27], for which there is bulk sequencing at two or more timepoints, we average the mutation rate calculated at each of these timepoints. (79) is applied for each timepoint with the respective CCFs and observed *γ* values for each timepoint.

The number of passengers present in the original type *i* founder cell cannot be directly observed, but we can measure *m*_*i*_, the number of clonal passengers present in the type *i* population, only including passengers not present in other clones. We will assume that the *m*_*i*_ don’t contain any shared mutations, which is true for the CLL dataset we consider. The likelihood function *P* (*m*_1_, …, *m*_*k*_|*t*_1_, …, *t*_*k*_) is proportional to

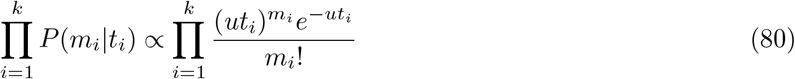

Then, maximizing the logarithm of (80) with respect to *t*_1_, *t*_2_, …, *t*_*k*_,

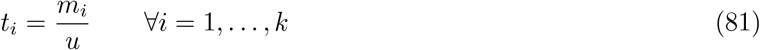

The observed clonal passengers in the founder type-*i* cell will incorrectly include passengers that reached fixation in the type-*i* population after driver mutation *i* occurred, instead of correctly being counted toward the descendant of clone *i*. As a result, we again correct for the expected number of these passengers, *u/r*_*i*_. That is,

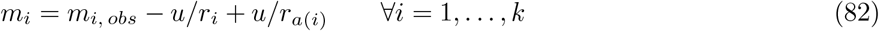

## Availability of data and materials

All simulated data generated during this study are included in this published article and its supplementary information files. CLL data analyzed is publicly available in Supplementary Tables from Ref. [27]. Code can be found at https://github.com/nathanlee543/Cancer_Inf_Sims

