## Supplementary materials for "Inferring parameters of cancer evolution from sequencing and clinical data"

#### Supplementary Table

Table 1: Inferred parameters for CLL patients with exponential growth patterns, for which there are at least two longitudinal bulk sequencing measurements before treatment. Estimates are computed from tumor size measurements and mutation frequencies from whole exome sequencing. Mutation rates are for the exome only. The time estimates are in terms of the patient's age in years. $f_1 = 0.01$, $f_2 = 0.2$, $L = 2$.

| Parameter | Pt. 3 | Pt. 6 | Pt. 9 | Pt. 21 |
|---|---|---|---|---|
| $r$ (/yr) | 0.505 [0.199, 0.847] | 0.676 [0.153, 1.295] | 0.279 [0.165, 0.416] | 0.793 [0.302, 1.141] |
| $r_1$ (/yr) | 0.852 [0.652, 1.042] | 0.407 [0.08, 0.731] | -0.396 [-0.453, -0.193] | 1.518 [1.007, 2.037] |
| $r_2$ (/yr) | | 0.458 [0.084, 0.846] | 0.672 [0.487, 0.94] | |
| $r_3$ (/yr) | | 1.087 [0.65, 1.777] | 0.626 [0.388, 0.863] | |
| $u$ (mut/yr) | 0.479 [0.394, 0.589] | 0.149 [0.123, 0.194] | 0.361 [0.349, 0.372] | 0.204 [0.191, 0.228] |
| MRCA (yr) | 14.6 [1.4, 26.8] | 2.8 [0.1, 13.2] | 4.9 [1.2, 10.8] | 6.4 [0.3, 16.7] |
| $t_1$ (yr) | 33.5 [24.1, 39.2] | 35.4 [21.7, 46.1] | 18.8 [8.8, 35.1] | 19.6 [10.8, 24] |
| $t_2$ (yr) | | 46.7 [25.6, 57.5] | 21.3 [7.7, 31.7] | |
| $t_3$ (yr) | | 45.9 [31.3, 54.6] | 24.8 [10.3, 37.6] | |

### Supplementary Figures

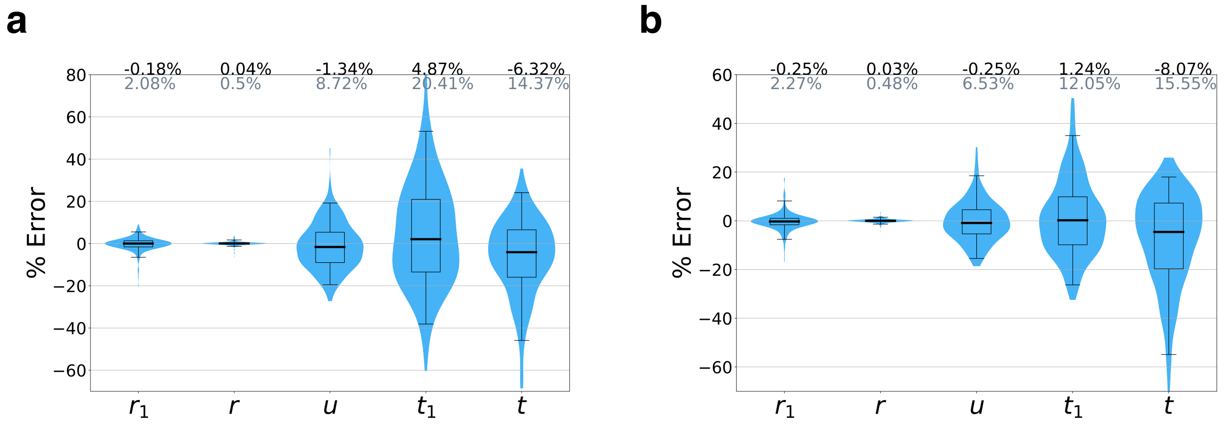


Figure S1: **Percent errors (PEs) for case with no death**. Accuracy of parameter inferences for surviving Monte Carlo simulation runs with ground truth parameter set: $b = 0.25$, $b_1 = 0.375$, $d = d_1 = 0.0$, $t_1 = 23$, $t = 17$, $\delta = 6$, $f_1 = 1\%$, and $f_2 = 20\%$. Mean percent error (MPEs) are the black numbers above the plots, and mean absolute percent errors (MAPEs) are the grey numbers below the MPEs. Boxes contain 25th-75th quartiles, with median indicated by thick horizontal black line. Whiskers of boxplots indicate 2.5 and 97.5 percentiles. Violins are smoothed density estimates of the percent error datapoints. (a) mutation rate $u = 1$. 290 surviving runs. (b) $u = 10$. 301 surviving runs.

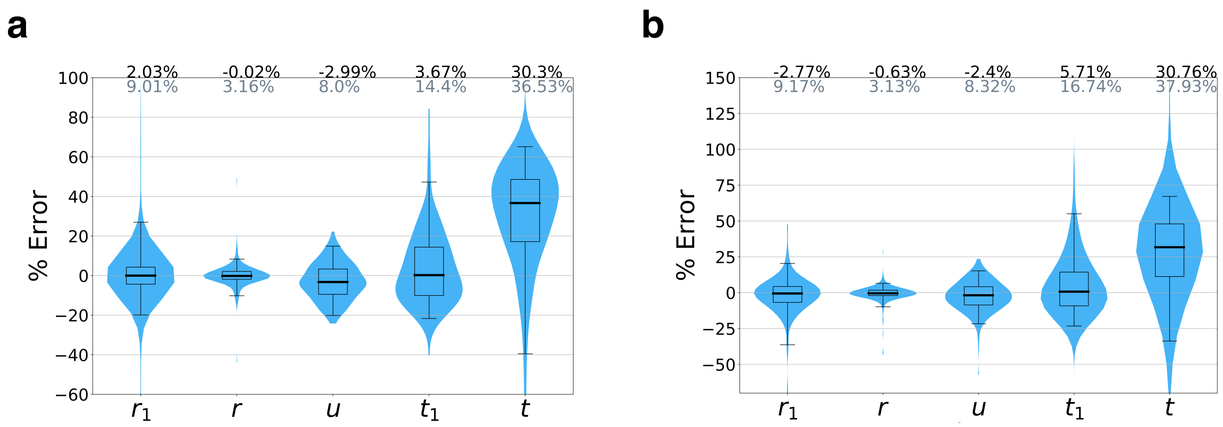


Figure S2: **Percent errors (PEs) for slow growing tumor**. Accuracy of parameter inferences for surviving Monte Carlo simulation runs with ground truth parameter set: $b = 0.25$, $b_1 = 0.25$, $d = 0.225$, $d_1 = 0.2125$, $t_1 = 180$, $t = 135$, $\delta = 45$, $f_1 = 1\%$, and $f_2 = 20\%$. Mean percent error (MPEs) are the black numbers above the plots, and mean absolute percent errors (MAPEs) are the grey numbers below the MPEs. Boxes contain 25th-75th quartiles, with median indicated by thick horizontal black line. Whiskers of boxplots indicate 2.5 and 97.5 percentiles. Violins are smoothed density estimates of the percent error data points. (a) mutation rate $u = 1$. 225 surviving runs. (b) $u = 5$. 249 surviving runs.

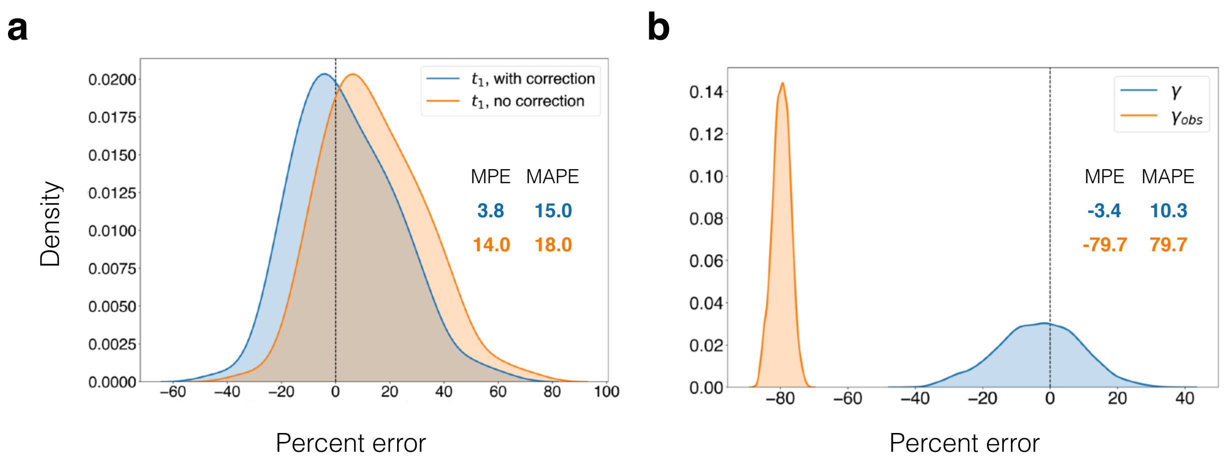


Figure S3: **Corrections for observed mutation counts.** (a) We compare percent errors of parameter estimates for time from tumor initiating until appearance of a driver subclone, $t_1$, with and without the correction for passengers that occur after the driver and reach fixation in the driver population (Eq. (6), main text). Errors for estimate with correction (Eq. (12), main text) are shown in blue, and for estimate without correction (Eq. (5), main text) in orange. Errors are plotted as a kernel density estimate for Monte Carlo simulations of fast growing tumor with mutation rate $u = 1$. Mean percent errors (MPEs) and mean absolute percent errors (MAPEs) are listed. (b) The percent errors for the observed (orange) and corrected (blue) number of subclonal mutations between frequencies $f_1$ and $f_2$, $\gamma$, (Eq. (7), main text) are plotted as kernel density estimates. Observed mutations are those that passed post-processing, i.e. those that have more than $L = 2$ mutant reads. True mutation frequencies were generated from 135 surviving runs of a Monte Carlo simulation of a fast growing tumor with mutation rate $u = 1$, from which sequencing reads were simulated with 100x average coverage (see Materials and Methods). Percent errors are calculated relative to the true $\gamma$ measured from the true mutation frequencies.

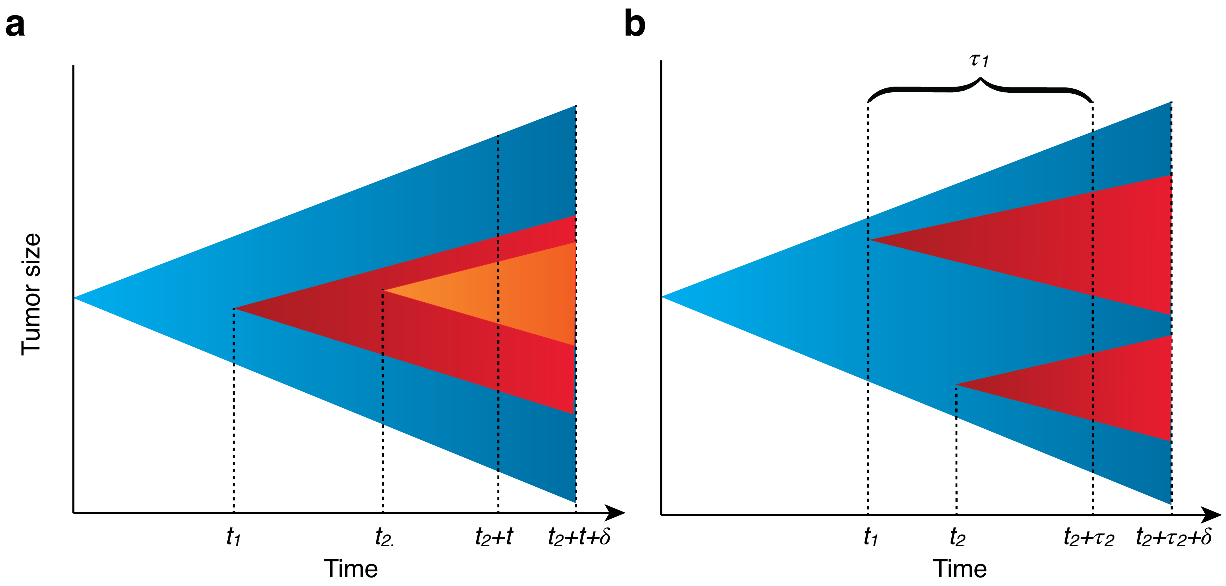


Figure S4: **Model for tumor expansion with two driver mutations.** (a) Two nested driver subclones. Initiated tumor (type-0) cells in blue, cells with driver 1 (type-1) in red, and cells with both drivers (type-2) in orange. A driver mutation occurs in a type-0 cell at $t_1$. A second driver mutation occurs in a type-1 cell at $t_1 + t_2'$. Tumor is bulk sequenced at $t_1 + t_2' + t$ and $t_1 + t_2' + t + \delta$. (b) Two sibling driver subclones. Type-0 cells (in blue). A driver mutation occurs in a type-0 cell at $t_1$. A second driver mutation occurs in a different type-0 cell at $t_2$. Tumor is bulk sequenced at $t_1 + \tau_1$ (or, equivalently $t_2 + \tau_2$) and $t_1 + \tau_1 + \delta$ (equivalently $t_2 + \tau_2 + \delta$).
